# Substrate-dependent accessibility changes in the Na^+^-dependent C_4_-dicarboxylate transporter, VcINDY, suggest differential substrate effects in a multistep mechanism

**DOI:** 10.1101/2020.01.16.902445

**Authors:** Connor D. D. Sampson, Matthew J. Stewart, Joseph A. Mindell, Christopher Mulligan

**Affiliations:** School of Biosciences, University of Kent, Canterbury, Kent, CT2 7NH, United Kingdom; Membrane Transport Biophysics Section, Porter Neuroscience Research Center, National Institute of Neurological Disorders and Stroke, National Institutes of Health, Bethesda, MD 20892, USA

## Abstract

Members of the <u>d</u>ivalent <u>a</u>nion <u>s</u>odium <u>s</u>ymporter (DASS) family (SLC13 in humans) play critical roles in metabolic homeostasis, influencing many processes including fatty acid synthesis, insulin resistance, adiposity, and lifespan determination. DASS transporters catalyse the Na^+^-driven concentrative uptake of Krebs cycle intermediates and sulfate into cells; disrupting their function can protect against age-related metabolic diseases and can extend lifespan. An inward-facing crystal structure and an outward-facing model of a bacterial DASS family member, VcINDY from *Vibrio cholerae*, predict an elevator-like transport mechanism involving a large rigid body movement of the substrate binding site. How substrate binding influences the conformational state of VcINDY is currently unknown. Here, we probe the interaction between substrate binding and VcINDY conformation using a site-specific alkylation strategy to probe the solvent accessibility of several broadly distributed positions in VcINDY in the presence and absence of substrates (Na^+^ and succinate). Our findings reveal that accessibility to all positions tested can be modulated by the presence of substrates, with the majority becoming less accessible in the presence of Na^+^. Mapping these solvent accessibility changes onto the known structures of VcINDY suggests that Na^+^ binding drives the transporter into an as-yet-unidentified intermediate state. We also observe substantial, separable effects of Na^+^ and succinate binding at several amino acid positions suggesting distinct effects of the two substrates. Furthermore, analysis of a solely succinate-sensitive residue indicates that VcINDY binds its substrate with a low affinity and proceeds via an ordered process in which one or more Na^+^ ions must bind prior to succinate. These findings provide insight into the mechanism of VcINDY, which is currently the only structural-characterised representative of the entire DASS family.

## Introduction

The divalent anion sodium symporter (DASS) family of transporters are present in all domains of life and are responsible for the transport of several key compounds into cells, such as citrate, Krebs cycle intermediates and sulfate^1^. Cytoplasmic citrate plays a major role in the metabolism of eukaryotic cells, contributing to the regulation of fatty acid, cholesterol and low-density lipoprotein synthesis^2-5^. By maintaining and controlling the cytoplasmic citrate concentration, members of the DASS family (Transport Classification Database no. 2.A.47, SLC13 in humans) are key players in metabolic regulation in eukaryotes as demonstrated by the phenotypes associated with their functional disruption. Knockdown of a DASS family member in fruitflies and nematodes leads to phenotypes analogous to caloric restriction, most notably a substantial increase in the lifespan of the organism, hence the alternative name for this family, INDY, which stands for *I’m not dead yet*^6^. In mice, knockout of a DASS family member (NaCT, SLC13A5 in humans) leads to protection against adiposity and insulin resistance^7^, and knockout of the equivalent transporter in human hepatocarcinoma cells substantially reduced hepatoma cell proliferation and colony formation^8^. In addition to metabolic diseases, mutations in a DASS family member (NaCT, SLC13A5) are associated with neurological disorders including developmental delay and early onset epilepsy^9^. Thus, DASS family members are prime targets for therapeutics designed to tackle metabolic diseases including diabetes and obesity, liver cancer and neurological disorders.

DASS transporters are ion-coupled secondary active transporters. Secondary transporters can harness the energy stored in ion gradients (usually Na^+^ or H^+^) across the membrane to drive the energetically uphill movement of substrate against its concentration gradient. Secondary active transporters occupy at least two major conformational states, the inward-facing state (IFS) and the outward-facing state (OFS), which alternate in order to expose the substrate binding site from the cytoplasmic to the extracellular side of the membrane, and vice versa.

The DASS transporter family belongs to the Ion Transporter (IT) Superfamily^10,11^, and the majority of characterised DASS transporters are known to transport their anionic substrates coupled to the co-transport of multiple Na^+^ ions^12 13-17^. All of our current structural and mechanistic understanding of the DASS family comes from studies on a bacterial family member, VcINDY, from *Vibrio cholerae*, which is the only DASS transporter for which we have high resolution structural information^18,19^. Functional characterisation of VcINDY reveals it preferentially transports C_4_-dicarboxylates, for example, succinate, malate and fumarate^17^. The model VcINDY substrate, succinate, is transported in its dianionic form coupled to the transport of three Na^+^ ion^17,20^. Our lab and others have revealed that VcINDY shares structural and functional characteristics with its mammalian homologues, suggesting that insight derived from the mechanism of VcINDY is directly applicable to the mammalian transporters^17,20,21^.

The x-ray structures of VcINDY reveal that it forms a homodimer and each protomer consists of two domains; the scaffold domain that forms dimer interface contacts, and the transport domain that houses key substrate binding residue (Fig. 1A)^18,19^. All structures of VcINDY to date are captured in the same IFS conformation where the substrate is exposed to the cytoplasmic side of the membrane suggesting that the crystallisation conditions select for this lowest free-energy state conformation of VcINDY^18,19^. Using a combination of symmetry-based structural modelling and extensive biochemical and biophysical validation, we have described an OFS conformation of VcINDY in which the transport domain and the substrate binding site are vertically translocated through the membrane (Fig. 1A)^22^. Combined, the IFS structure and OFS model predict that VcINDY employs an elevator-like mechanism to achieve alternating access to the substrate binding site from both sides of the membrane.

**Figure 1.**
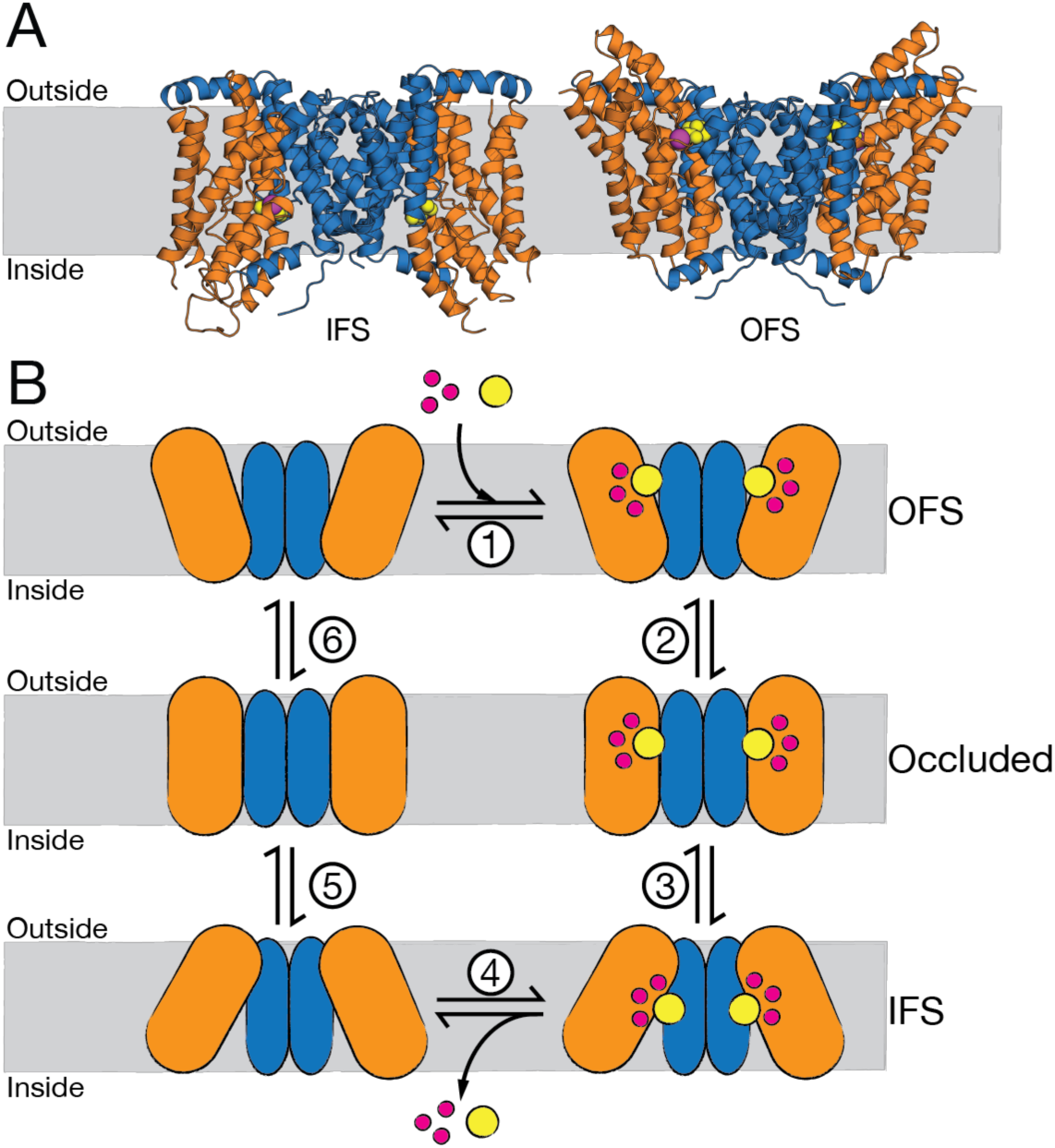
Structure and simple kinetic transport scheme of VcINDY. **A)** Inward-facing state (IFS, left) crystal structure (PDB: 4F35) and outward-facing state (OFS, right) model of VcINDY. Transport domain is depicted as orange helices and the scaffold domain is blue. Bound substrate is yellow spheres; the bound Na^+^ ion is a magenta ball; membrane is indicated by the grey rectangle. **B)** Simple kinetic model of transport by VcINDY. Substrate-free OFS (top left) binds Na^+^ and succinate^2-^ in an unknown order (step 1), at which point the transporter transitions from OFS to IFS via an occluded state (steps 2 & 3). Substrates are released into the inside of the cell (step 4), and the empty transporter undergoes a conformational change back to the substrate-free OFS (steps 5 & 6). Colour scheme is the same as (A).

The structures of VcINDY reveal that the substrate binding site within the transport domain is composed of backbone and side chain contacts from the tips of two re-entrant hairpin loops and an unwound helix (TM 7)^18^. This organisation is reminiscent of other elevator-like ion-driven transporters where local conformational changes of hairpin loops are required for ligand binding, controlling coupling ion and substrate access to the binding site, and as a means of preventing uncoupled transport^23,24^. The transport of substrate and coupling ion is tightly coupled, meaning that Na^+^ transport cannot occur without succinate transport, and vice versa. In a simple kinetic scheme for VcINDY; succinate^2-^ and 3 Na^+^ ions bind to the OFS (Fig 1 B, step 1), facilitating a conformational change into the IFS via an intermediate occluded state that is yet to be structurally characterised (Fig 1 B, step 2, 3). The substrates are released from the IFS (Fig 1 B, step 4), and the empty transporter transitions back to the OFS via an occluded intermediate state (Fig 1 B, step 5, 6). The key to tightly coupled transport is that the IFS-to-OFS transition *cannot* occur unless the transporter either is ligand-free or carrying its full complement of substrate and coupling ions. What remains to be determined is how the presence of substrates influences the conformational state of VcINDY and whether local conformational changes of the re-entrant hairpins are required for transport.

Site-directed alkylation of single cysteine residues has been a valuable tool in elucidating the dynamic features of transporters, in particular the pioneering work on lactose permease by Kaback and co-workers^25,26^. Alkylation of single cysteine residues provides a readout of the accessibility of a particular position on the protein. The reactivity of single cysteine residues to hydrophilic thiol-reactive reagents depends on the solvent accessibility of the amino acid residue in a given conformation. Thus, any change in the reactivity between the cysteine and the thiol-reactive reagent reflects a change in the local environment and/or solvent accessibility to that particular position in the protein.

Here, we explore the substrate dependence of the elevator-like conformational changes of VcINDY. To achieve this, we employ the hydrophilic, thiol-reactive reagent, methoxypolyethylene glycol maleimide 5K (mPEG5K) to probe the solvent accessibility of substituted single cysteines that are predicted to be accessible in the IFS or the OFS, but not both, according to the IFS crystal structure and OFS model. Our findings are consistent with VcINDY entering a structurally uncharacterised intermediate conformation upon binding substrates, with the majority of these accessibility changes being induced by binding Na^+^ alone. Several positions towards the tips of the hairpin loops were identified that, unlike all other positions tested, displayed substantial sensitivity to the binding of succinate over and above the sensitivity to Na^+^ binding. Taken together, these observations are consistent with a transport mechanism that involves multiple conformational changes that differ in their substrate dependence. In addition, our findings suggest that Na^+^ must bind prior to succinate and that dicarboxylate substrates appear to bind with a very low affinity.

## RESULTS

### Experimental approach and generation of substituted cysteine panel

In this work, we sought to determine whether the presence of substrates drives VcINDY predominantly into its IFS, its OFS, or into an intermediate conformational state that has not yet been structurally characterised. To probe the conformational state of VcINDY, we devised a substituted cysteine solvent accessibility assay in which we could measure the rate of modification of substituted cysteine residues using a hydrophilic cysteine reactive mass tag, mPEG5K, in the presence and absence of substrates (Fig. 2A). The reaction between an accessible single cysteine residue and mPEG5K would result in a 5 kDa increase in protein mass that is discernible on SDS-PAGE due to the PEGylated protein’s retarded electrophoretic mobility (Fig. 2B). Digitisation and densitometric analysis of the distribution of bands in each sample allows us to quantify the extent of PEGylation and calculate a modification efficiency for each timepoint (Fig. 2C). Plotting the modification efficiency for each sample in a timecourse provides us with a means of monitoring the modification rate of a particular position on the protein.

**Figure 2.**
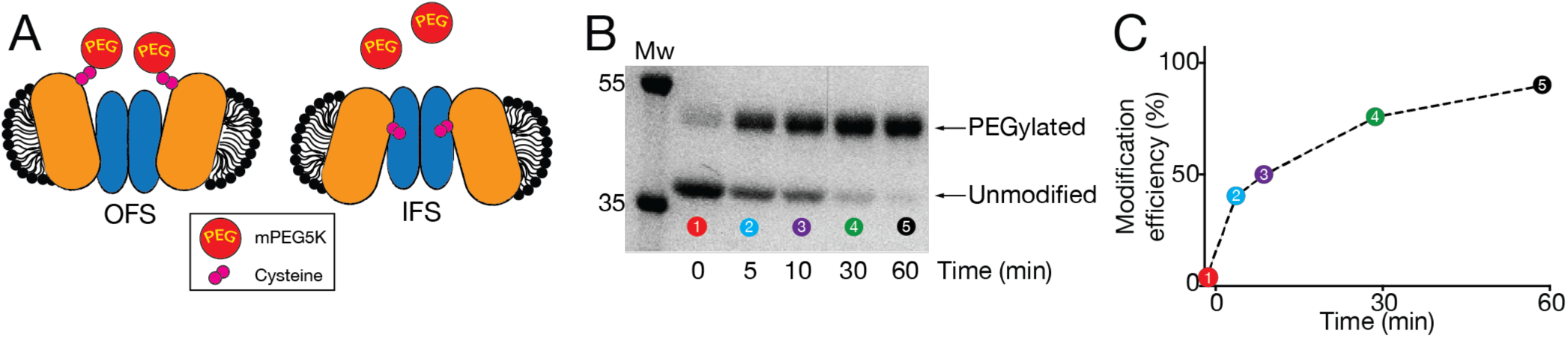
Probing solvent accessibility of substituted cysteines. **A)** Cartoon representation of VcINDY in its outward-facing state (OFS, left) and inward-facing state (IFS, right) depicting the change in solvent accessibility of a single substituted cysteine (pink spheres) in these different conformations. Cysteines that are more solvent exposed will react more readily with the mPEG5K (red spheres) resulting in increased rate of PEGylation. VcINDY’s colour scheme is the same as Figure 1. **B)** An SDS-PAGE gel depicting the gradual modification of a single cysteine residue of VcINDY by mPEG5K. Upon modification with mPEG5K, the apparent molecular weight of VcINDY increases from ∼38 kDa to 43 kDa. Band intensity is quantified by densitometry. **C)** Graph of the modification efficiency of the VcINDY cysteine mutant as a function of time using data derived from the SDS-PAGE gel shown in (B).

To allow us to differentiate between the IFS and OFS, we selected residues predicted to be accessible in the IFS or the OFS, but not both, using the IFS crystal structure and OFS repeat-swapped model as guides^18,22^. While avoiding highly conserved residues or residues involved in substrate or cation binding, we selected 24 amino acids in VcINDY to be individually substituted for cysteine, using a functionally active cysteine-free version of VcINDY as a background (see Table 1 for a full list of mutants tested)^17^. Of the 24 single cysteine variants produced, we discarded those mutants that were not produced in sufficient quantities for analysis, were incapable of catalysing Na^+^-driven succinate transport, or were not reactive to mPEG5K under any conditions (Table 1). Following this sieving step, we were left with 8 single cysteine mutants; VcINDYA120C^OFS^, VcINDYT215C^OFS^, VcINDYS381C^OFS^, VcINDYL384C^OFS^, VcINDYV388C^OFS^ that are predicted to be more accessible in the OFS (^OFS^ denoting that they are predicted to be OFS-accessible cysteines); VcINDYT154C^IFS^, VcINDYM157C^IFS^, and VcINDYT177C^IFS^ that are predicted to be more accessible in the IFS (^IFS^ denoting their predicted IFS accessibility); and a control cysteine mutant, VcINDYE42C, which is predicted to be equally accessible in both IFS and OFS. Importantly, all of the single cysteine mutants used in this analysis were capable of catalysing Na^+^-drive succinate transport, demonstrating that they can sample all transport-relevant conformations (Supplementary Figure 1). Using this panel of substituted cysteine mutants and the experimental approach described above, we sought to determine whether the presence or absence of substrates (Na^+^ and succinate) drives VcINDY into the IFS, OFS, or a hitherto unidentified intermediate.

**Table 1.** List of VcINDY cysteine variants produced during the course of this work. For each variant, it is indicated whether the protein could be stably expressed, whether it was active in *in vitro* transport assays, whether it was modifiable with mPEG5K, and if so, could this modification be modulated by substrate.

| Cysteine mutant | Expressed? | Active? | Modifiable by mPEG5K? | Modulated by substrate? |
| --- | --- | --- | --- | --- |
| E42C | Yes | Yes | Yes | No |
| F79C | Yes | Yes | No | - |
| A120C | Yes | Yes | Yes | Yes |
| W148C | Yes | Yes | No | - |
| T154C | Yes | Yes | Yes | Yes |
| M157C | Yes | Yes | Yes | Yes |
| T177C | Yes | Yes | Yes | Yes |
| Y178C | Yes | Yes | No | - |
| A206C | No | - | - | - |
| A208C | Yes | Yes | No | - |
| S213C | No | - | - | - |
| T215C | Yes | Yes | Yes | Yes |
| V272C | Yes | Yes | No | - |
| A346C | Yes | Yes | No | - |
| V364C | Yes | Yes | No | - |
| V370C | Yes | No | - | - |
| F375C | Yes | No | - | - |
| S381C | Yes | Yes | Yes | Yes |
| A383C | Yes | No | - | - |
| A384C | Yes | Yes | Yes | Yes |
| V388C | Yes | Yes | Yes | Yes |
| T391C | No | - | - | - |
| G430C | Yes | Yes | No | - |
| S436C | Yes | Yes | No | - |

### VcINDY likely favours an intermediate conformation in the presence of saturating substrate concentrations

To provide us with a readout on the conformational state of VcINDY, we selected two mutants for initial analysis that, based on available structural information, we predict are only accessible in the IFS or the OFS; VcINDYT154C^IFS^ (Fig 3A, left) and VcINDYS381C^OFS^ (Fig. 3A, middle), respectively. To assess substrate-induced changes to the modification efficiency of these positions, we incubated each mutant protein with saturating substrate concentrations (1 mM succinate, 150 mM NaCl) or under apo conditions (no succinate and with Na^+^ ions replaced with experimentally inert K^+^), and quantified the rates of modification using SDS-PAGE and densitometric analysis (Fig. 3 B and C).

**Figure 3.**
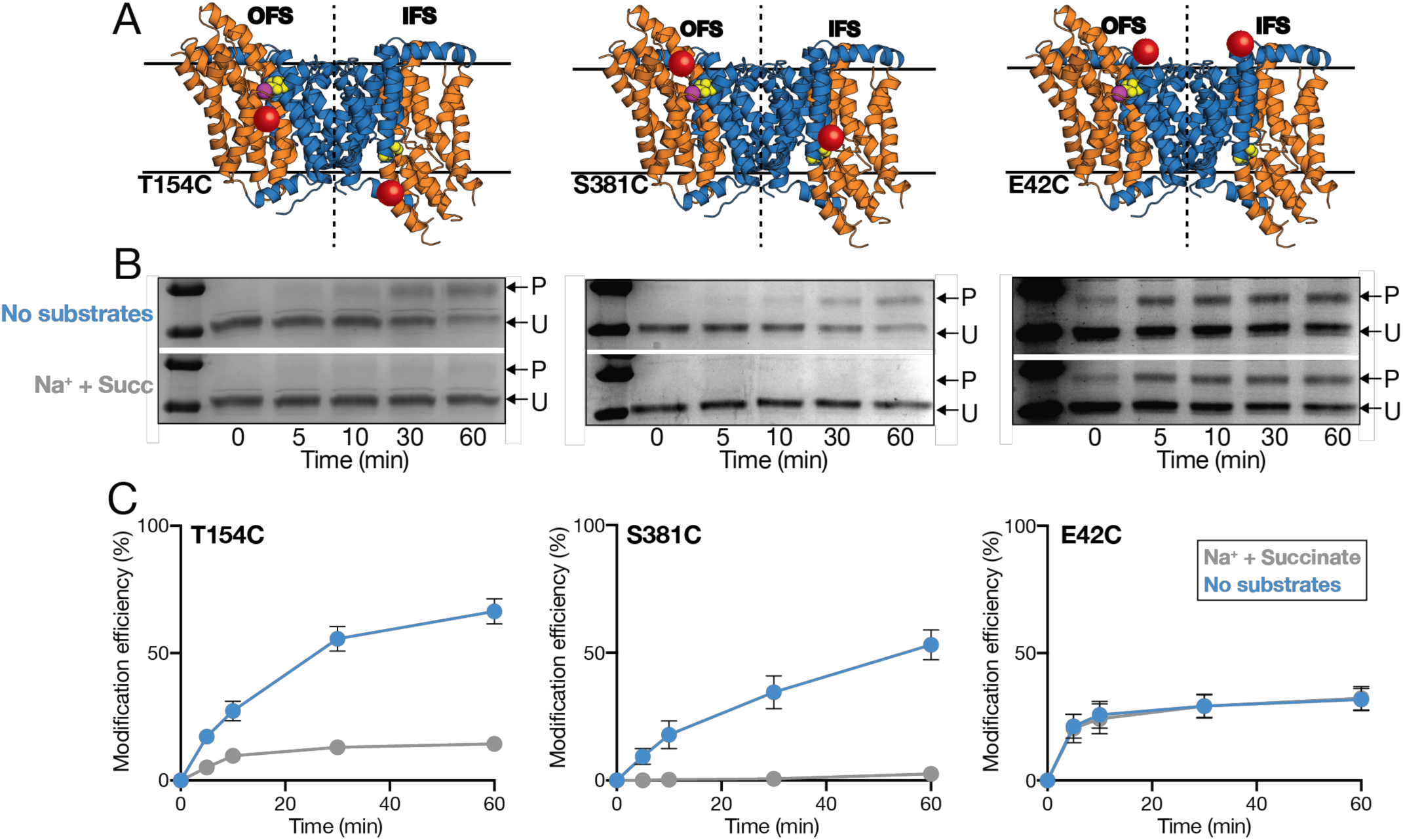
Substrate-induced accessibility changes in VcINDY. **A)** A merge of the structures of the outward-facing state (OFS, left of the dotted line) and the inward-facing state (IFS, right of the dotted line) illustrating the relative accessibility of the single substituted cysteine residues (red spheres); T154C (left panel), S381C (middle panel), and E42C (right panel) in each conformation. The colour scheme used for VcINDY is the same as in Figure 1. **B)** Representative SDS-PAGE gels of PEGylation timecourse of the single cysteine mutants T154C (left panel), S381C (middle panel), and E42C (right panel) in the presence (bottom gel) and absence of substrates (top gel). The PEGylated protein bands (P) and unmodified protein bands (U) are indicated by arrows. **C)** Proportion (%) of each single cysteine mutant modified at each timepoint in the presence (grey data) of saturating Na^+^ and succinate and in apo conditions (blue data). Data points are the average of triplicate datasets and the error bars represent SEM. This experiment was performed on at 3 separate occasions with the same result.

In the apo state, we observe substantial PEGylation of the IFS-accessible VcINDYT154C^IFS^ over the timecourse of the experiment resulting in PEGylation of ∼65% of the protein (Fig. 3C, left, blue data). However, in the presence of saturating substrates, PEGylation of this position was almost completely prevented, suggesting that VcINDY favours a conformation in which T154 is not solvent accessible under these conditions (Fig. 3C, left, grey data).

Based on our current understanding of the structural mechanism of VcINDY, these data suggest that the presence of substrates drives VcINDY into a non-IFS conformation, which, based on our 2-state model, is the OFS. We reasoned that if this change in accessibility of T154C is indeed due to a rigid body elevator-like movement of VcINDY’s transport domain, then we should observe the opposite effect of substrates on a single cysteine in a position predicted to be accessible in the OFS, but not in the IFS. To test this hypothesis, we analysed the PEGylation rate of the OFS-accessible cysteine variant VcINDYS381C^OFS^, in the presence and absence of substrates. In the absence of substrates, we observed a steady labelling rate of VcINDYS381C^OFS^ over the course of our experiment (Fig. 3B and C, middle panel). However, instead of seeing a further increase in the modification rate upon addition of substrate, which would be consistent with our hypothesis, we observe that labelling of VcINDYS381C^OFS^ is almost completely blocked in the presence of Na^+^ and succinate (Fig. 3B and C, middle panel).

To rule out the possibility that the presence of substrate could be directly diminishing the reactivity of mPEG5K through some unforeseen mechanism, we performed our PEGylation assay on VcINDYE42C, whose single cysteine is predicted to be equally accessible in both of the known conformations of VcINDY (Fig. 3A, right). For this control mutant, we observed equal PEGylation rates in the presence and absence of substrates, indicating that the effect on the modification efficiency of our IFS- and OFS-accessible mutants is indeed due to interaction of the substrate with VcINDY (Fig. 3B and C, right panel).

Intriguingly, our data for VcINDYT154C^IFS^ and VcINDYS381C^OFS^ indicates that *both* positions become less accessible in the presence of substrates, which is consistent with VcINDY adopting an as-yet-unknown intermediate conformational state in the presence of substrates. To explore this possibility further, we analysed the modification rates of the 6 other VcINDY variants with single cysteines positioned in sites spanning the scaffold domain-transport domain interface; two predicted to be only accessible in the IFS, and 4 OFS-accessible only (Fig. 4A). Each of these 6 cysteine variants could be robustly PEGylated in the absence of substrates with final proportions of PEGylated protein ranging between 60-85%; these variations likely reflect the relative solvent accessibility of each position (Supplementary figures 2 and 3). While we also observed mutant-to-mutant variation in the magnitude of substrate-induced modification efficiency changes, *all* of the positions tested exhibited substantially reduced levels of modification in the presence of substrates compared to the absence, signified by a negative Δmodification efficiency value (Fig. 4B). These results indicate that all of the positions tested are less solvent accessible in the presence of substrates, further suggesting that VcINDY favours an intermediate state that is substantially different to the currently known conformations of VcINDY.

**Figure 4.**
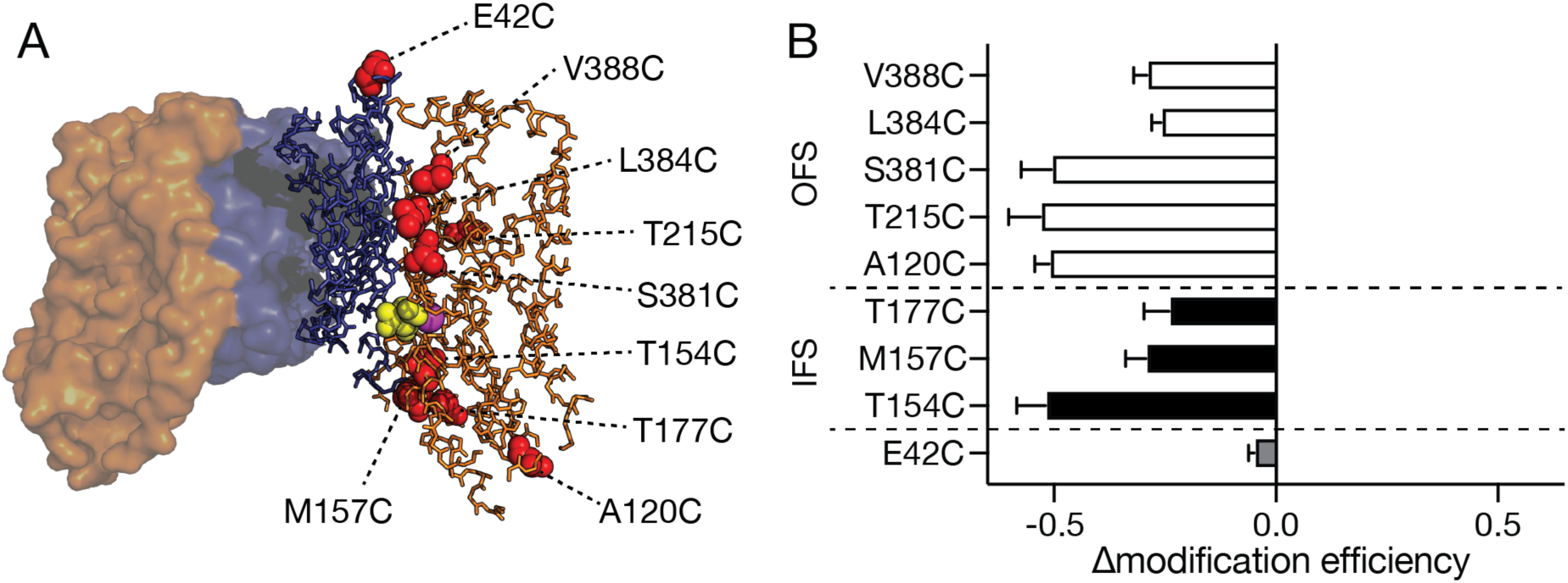
Summary of the effects of the addition of substrates (Na^+^ and succinate) on the accessibility of all single cysteine mutants. **A)** IFS X-ray structure of VcINDY showing the positions of each single cysteine substitution used in this analysis. Colour scheme is the same as Fig. 1. Each of the positions mutated to cysteine are shown in red spheres. **B)** Change in modification efficiency (Δmodification efficiency) is the difference between the modification efficiency at 60 min in the presence and absence substrates. Using the value obtained in the absence of substrates as a baseline, a negative value indicates lower modification efficiency in the presence of substrate; and a positive value indicates higher substrate-induced modification efficiency. Filled bars are the predicted IFS accessible mutants, the open bars are the predicted OFS-accessible mutants, and the grey bar is the control. The data are an average of at least 3 data sets and the error bars indicate SEM. This experiment was performed for each mutant at least 3 times.

### Modification rates are influenced by the presence of Na^+^

To investigate the substrate-induced changes in modification efficiency in more detail, we sought to determine the individual effects of the coupling ion, Na^+^, and the substrate, succinate, on the modification efficiency of each substituted cysteine. To do this, we measured the PEGylation rate of our panel of single cysteine mutants in one of 4 different conditions; Na^+^ alone; succinate alone; apo; or Na^+^ and succinate. In each case, we ensured that the reactions were osmotically and ionically balanced using KCl, which is known not to catalyse transport nor interact specifically with VcINDY^17,20^.

We first analysed the effects of coupling ion or substrate on the modification rates of the IFS- and OFS-accessible mutants, VcINDYT154C^IFS^ (Fig. 5A) and VcINDYS381C^OFS^ (Fig. 5B). For these two mutants, we observed no changes in the modification rate in the presence of succinate alone when compared to apo conditions (Fig. 5A and B). However, the presence of Na^+^ alone substantially reduced the modification rate of these mutants, accounting for almost all of the reduction in the modification rate we observed in the presence of both Na^+^ and succinate (Fig. 5A and B). These data suggest that the presence of Na^+^ alone is able to induce a shift in the protein’s conformation that obscures these positions and reduces modification efficiency, whereas succinate binding by itself contributes minimally. In addition, the observation that succinate alone is unable to influence the modification rate of these positions indicates that one or more Na^+^ ions must bind to VcINDY prior to succinate binding. We next tested the other members of our single cysteine panel to see if Na^+^ binding also reduced their modification rate. In total, Na^+^ binding alone accounted for the majority of substrate-induced modification rate changes in half of the single cysteine mutants, including; two IFS-accessible mutants, VcINDY154C^IFS^ and VcINDY177C^IFS^, and two OFS-accessible mutants, VcINDYA120C^OFS^, VcINDYS381C^OFS^ (Fig. 5C).

**Figure 5.**
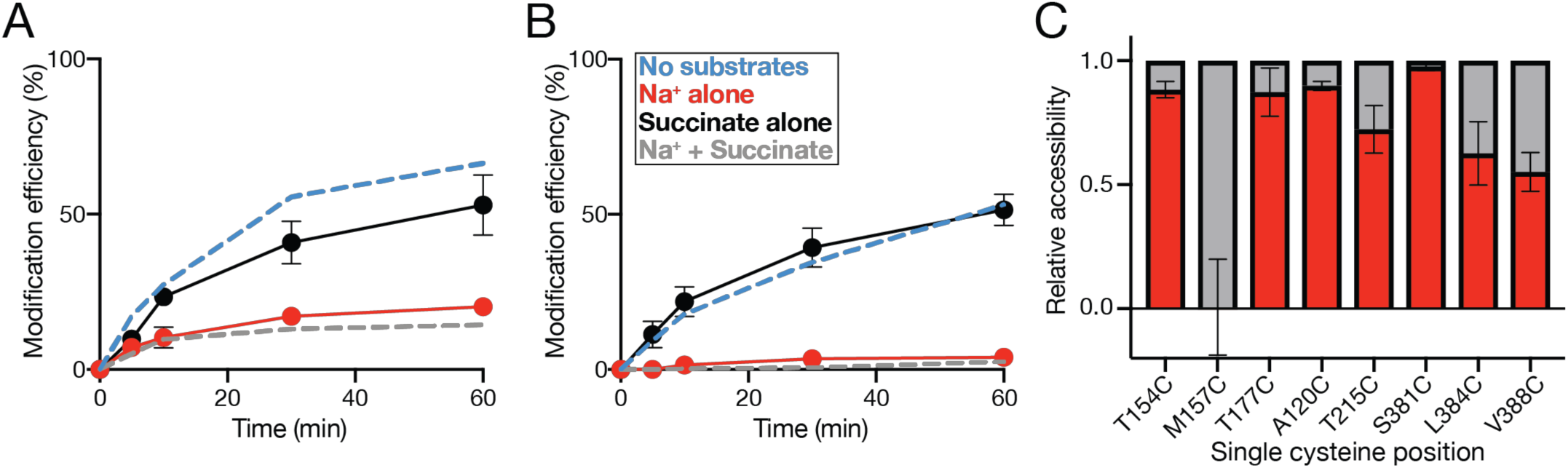
Effects of individual substrates on the single cysteine modification efficiency. Proportion (%) of **A)** VcINDYT154C and **B)** VcINDYS381C that is modified at each timepoint in the presence of Na^+^ alone (red data) or succinate alone (black data). Modification rate of each mutant in the presence and absence of saturating substrates (same data presented in Fig. 3) is shown as dashed lines for comparison (grey and blue data, respectively). **C)** Summary of the effects on the modification rates for each mutant of Na^+^ alone (red column) compared to the overall effect of Na^+^ and succinate (grey column). The data are an average of two data sets for M157C and 3 data sets all other mutants. Error bars indicate SEM. This experiment was performed for each mutant at least 3 times.

In addition to Na^+^-induced modification rate changes, 3 members of our mutant panel, VcINDY215C^OFS^, VcINDY384C^OFS^, and VcINDY388C^OFS^ also exhibited further substantial modification rate reduction upon addition of succinate in the presence of Na^+^. These additional succinate-induced modification efficiency changes raises the possibility that the sequential binding of each substrate stabilises discrete conformations of VcINDY.

Interestingly, the amino acid positions that exhibit separable sensitivity to both Na^+^ and succinate are located on the arms of the re-entrant hairpin loops that contribute to binding site formation and have been shown to perform a pivotal role in gating and coupling in other elevator-like transporters^23,24^. In contrast to all other positions tested, the modification efficiency of one of these re-entrant hairpin loop residues, VcINDYM157C^IFS^, was completely insensitive to the presence of Na^+^ ions alone, but exhibited substantial sensitivity to the presence of succinate (Fig. 5C and 6A). Succinate did not induce changes in the modification rate of VcINDYM157C^IFS^ in the absence of Na^+^, suggesting an ordered binding process in which one or more Na^+^ must bind prior to succinate binding (Fig. 5C and 6A).

While our data are consistent with the different substrate conditions stabilising particular conformations of VcINDY, an alternative possibility is that, in the presence of substrates, the fully loaded transport undergoes rapid IFS-OFS interconversion; so rapid perhaps that the maleimide, which has a relatively slow rate of reaction, does not have sufficient time to react, which would result in apparent inaccessibility. To test this possibility, we performed our PEGylation assay on VcINDYM157C^IFS^ in the presence and absence of succinate using MTS-PEG5K, whose methanethiosulfonate (MTS) moiety has a considerably faster reaction rate than maleimides^27^. Under these conditions, we observed the same decrease in modification efficiency in the presence of succinate compared to the absence (Supplementary Figure 4), indicating that our observations with mPEG5K reflect conformational stabilisation rather than altered protein dynamics.

### PEGylation rates of VcINDYM157C^IFS^ suggest ordered low affinity binding of succinate

Due to the maverick nature of the VcINDYM157C^IFS^ mutant, we investigated this variant in more detail. To determine whether the apparent succinate-induced decrease in modification efficiency was due to a specific interaction between VcINDY and succinate, we performed our PEGylation assay on VcINDYM157C^IFS^ in the presence of increasing concentrations of succinate ranging from no succinate up to 1 mM, while keeping a constant Na^+^ concentration of 150 mM (Fig 6 B).

**Figure 6.**
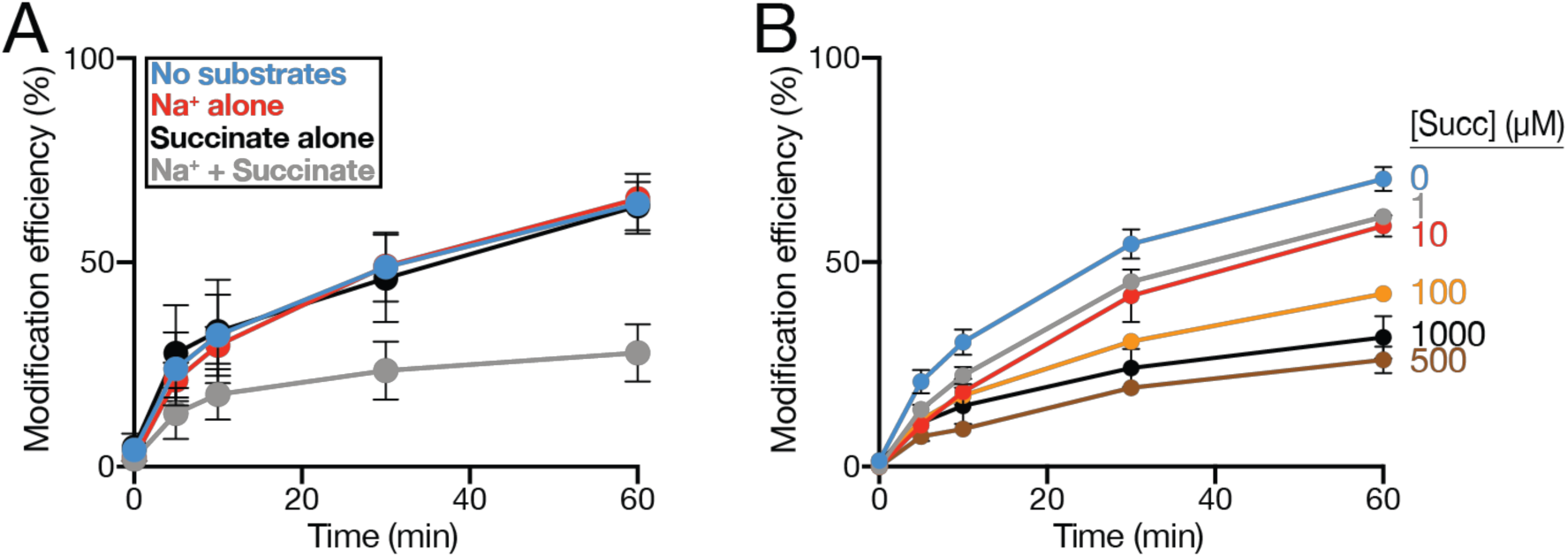
Effects of substrates on the modification efficiency of VcINDYM157C^IFS^. **A)** Modification efficiency (%) of VcINDYM157C^IFS^ at each timepoint in the presence of no substrate (150 mM K^+^, blue data), Na^+^ alone (red data), succinate alone (black data), and Na^+^ + succinate (grey data). **B)** Modification efficiency of VcINDYM157C^IFS^ in the presence of 150 mM NaCl and increasing concentrations of succinate from 0-1 mM. The data are an average of at least 3 data sets for A) and two data sets for B). Error bars indicate SEM. This experiment was performed for each mutant at least 3 times.

We observed a dose-responsive decrease in modification efficiency of VcINDYM157C^IFS^ with increasing succinate concentration, indicating that this effect is indeed due to succinate binding (Fig. 6B). However, substantial reduction in the modification efficiency is only apparent when the protein is incubated with ≥100 µM succinate, suggesting an unexpectedly low affinity for succinate, considering the *K*_m_ for transport is known to be 1 µM^17^.

In addition to succinate, VcINDY can transport a number of other C_4_-dicarboxylates, including malate, fumarate, and oxaloacetate, but not shorter dicarboxylates, such as oxalate, nor tricarboxylates, such as citrate^17^. If this effect on PEGylation is indeed due to VcINDY’s specific interaction with its substrates, we would expect other known substrates to be similarly influential, whereas non-transported dicarboxylates should have no effect. Compared to Na^+^ alone conditions, substituting succinate with malate in the PEGylation reaction resulted in a significant increase in the modification rate, whereas substituting succinate with the non-substrate oxalate resulted in modification rates akin to those observed in the absence of any dicarboxylate substrate (Fig. 7A). These data indicate that the reduction in modification rate of VcINDYM157C^IFS^ are due to specific substrate interactions and not due to unforeseen indirect effects of dicarboxylates. However, we noted with interest that the presence of malate led to substantially and significantly reduced protection of the cysteine from PEGylation compared to succinate (Fig. 7A).

**Figure 7.**
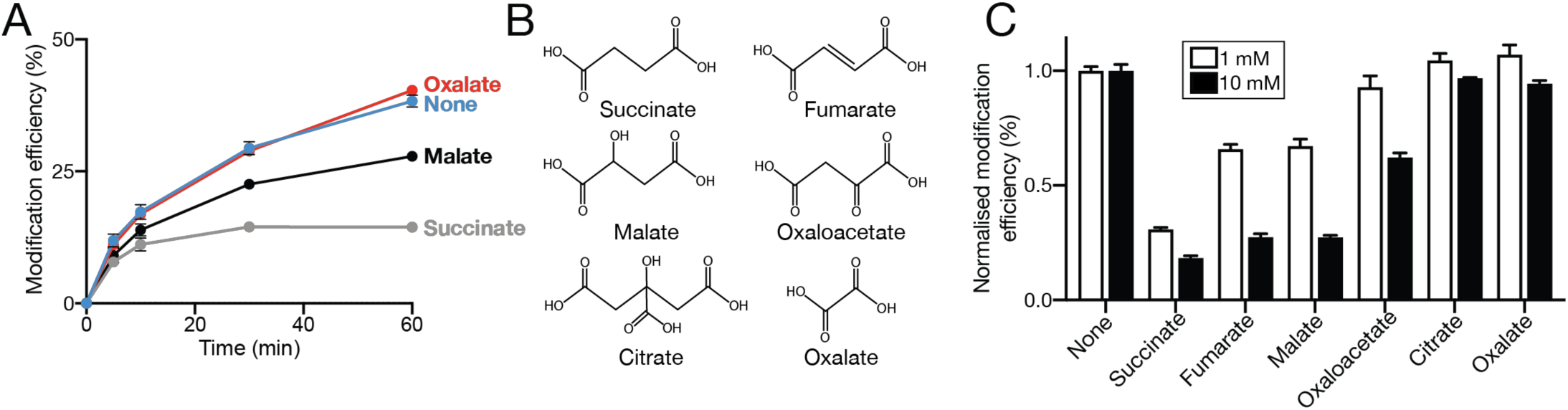
Effects of substrates on the modification efficiency of VcINDYM157C^IFS^. **A)** Proportion (%) of modified VcINDYM157C^IFS^ at each timepoint in the presence of 150 mM Na^+^ and no substrate (blue data), oxalate (red data), malate (black data), or succinate (grey data). **B)** Chemical structures of the compounds used in A) and B); succinate, fumarate, malate, oxaloacetate, citrate, oxalate. **C)** Normalised modification efficiency of VcINDYM157C^IFS^ after 1 hour in the presence of 150 mM NaCl and either 1 mM (open bars) or 10 mM (closed bars) of each indicated compound. The data are an average of at least 3 data sets and the error bars indicate SEM. This experiment was performed for each mutant at least 3 times.

Intrigued by this result, we extended the compound range in our VcINDYM157C^IFS^ PEGylation assay to include the other known VcINDY substrates, fumarate and oxaloacetate, and the non-substrate citrate (Fig. 7B). To test the effects of our extended compound range on the VcINDYM157C^IFS^ PEGylation rate, we determined the modification efficiency after 60 min incubation with mPEG5K and 1 mM of each test compound (Fig. 7C, open bars). As expected, the presence of succinate reduces the modification to the greatest extent, whereas oxalate, citrate, and surprisingly, oxaloacetate induce negligible changes to the modification efficiency compared to the absence of substrates (Fig. 7C). The presence of 1 mM fumarate and malate reduce modification efficiency substantially, but only half as much as succinate (Fig. 7C). We reasoned that these variations in the extent of labelling in the presence of different substrates could merely reflect differences in the affinity for each of the different substrates. To test this possibility, we performed the same PEGylation assay, but with a final concentration of 10 mM of each compound (Fig. 7C, closed bars). In the presence of increased concentrations of each compound, we observe no change in the modification efficiency in the presence of either oxalate nor citrate (Fig. 7C). Increasing the concentration of oxaloacetate to 10 mM results in significantly decreased modification efficiency compared to 1 mM, and while increasing the concentration of fumarate and malate to 10 mM leads to further reduction in the modification efficiency, they were still significantly less effective than 10 mM succinate (Fig. 7C). These data suggest that the differences in the modification efficiency of VcINDYM157C^IFS^ in the presence of different substrates is due primarily to differences in VcINDY’s binding affinity for each substrate. In addition, these results reveal that VcINDY’s substrate binding site is not saturated in the presence of 1 mM substrate, suggesting that under these experimental conditions, VcINDY has a remarkably low affinity for its substrates.

## DISCUSSION

In this work, we have described the first foray into determining the substrate-dependent conformational changes of VcINDY, which is a structural and mechanistic representative of the entire DASS transporter family. Using a site-specific alkylation strategy on detergent-solubilised protein, we have demonstrated that the modification rate, which serves as a proxy for solvent accessibility, of several broadly distributed positions in VcINDY can be modulated by the presence of substrates. We demonstrate that the majority of these changes in modification efficiency can be induced by the presence of Na^+^ alone. However, we also observe substantial, separable effects of succinate binding on the modification efficiency of multiple positions, suggesting discrete effects of the two substrates. Furthermore, we have identified a position whose modification rate is insensitive to the presence of Na^+^, but highly sensitive to succinate. Further analysis of this variant indicates that VcINDY binds its substrate with a low affinity and proceeds via an ordered process in which one or more Na^+^ ions must bind prior to the substrate, succinate. Mapping these solvent accessibility changes onto the IFS X-ray structure and OFS repeat-swapped model suggests that the binding of substrates drives the protein into an as-yet-unidentified intermediate state that obscures all of the positions tested.

### VcINDY likely undergoes multiple large- and small-scale conformational changes during transport

There are now many examples of structurally characterised transporters that are predicted to employ an elevator-like transport mechanism^24,28-37^, which has revealed common features amongst many of them, including; distinct scaffold and transport domains, a broken transmembrane helix containing an intramembrane loop that contributes to the substrate binding site, and two re-entrant hairpin loops that enter the membrane but do not fully span it. While structural differences exist between the predicted elevator-type transporters, they are all predicted (or have been shown) to undergo a substantial vertical translocation of the transport domain, usually accompanied by a pronounced rotation of this domain, to expose the substrate binding site to both sides of the membrane.

Our data suggest that multiple positions predicted to be solvent accessible in either the IFS or the OFS, but not both, of VcINDY become less accessible in the presence of Na^+^ (Fig. 4), consistent with the stabilisation of the an intermediate state of VcINDY. Cation-dependent conformational changes have been observed for multiple Na^+^-driven elevator-like transporters^38-40^. By occupying an intermediate state between the IFS and OFS, many of the residues tested could be obscured, leading to reduced modification. No intermediate state structure exists for VcINDY, so we cannot map the required conformational changes directly onto a structure of VcINDY. However, intermediate states of other elevator-like transporters, either crosslink-stabilised or captured in the presence of substrate^32,41^, have been structurally characterised, demonstrating that an intermediate state is well-occupied during the elevator-like mechanism. Indeed, in the case of the best-characterised elevator transporter, Glt_Ph_, the protein only transitions between the OFS and intermediate state, or the IFS and intermediate state in the presence of Na^+^ alone^40^. Only when both the cation and substrate are present can the protein fully transition between the IFS and OFS (via the intermediate state) ^40^. This blockage of the cation only-bound state is crucial to tight coupling in secondary active transporters and prevents cation leak, which could prove catastrophic to the cell.

In both VcINDY and Glt_Ph_, the tips of these re-entrant loops form part of the substrate binding site, coordinating both the coupling ion and substrate^18,19,42^. However, a major mechanistic difference between VcINDY and Glt_Ph_ is the role of the re-entrant loops in gating and coupling. In Glt_Ph_ and other glutamate transport homologues, the substrate and coupling ions are fully enclosed in the transport domain^23,42-44^. An OFS crystal structure of Glt_Ph_ in the presence of the bulky inhibitor D,L-threo-β-benzyloxyaspartate (TBOA) revealed that the outermost re-entrant loop, HP2, could be propped open to allow substrate access to the binding site^23^. The local and relatively subtle conformational changes of this hairpin are crucial to tight coupling of these transporters; the re-entrant loop is “open” in the absence of the full complement of substrate and coupling ions, which prevents premature translocation of the binding site that would lead to uncoupled transport. The symmetrically related re-entrant loop, HP1, was thought to play a similar role on the cytoplasmic face of the transporter. However, recent structural analysis of the human neutral amino acid transporter (ASCT2), which is structurally related to Glt_Ph_, revealed that HP2 is also required to open in the IFS; whereas HP1 remains static^24^.

In the IFS structure and OFS model of VcINDY, the substrate is solvent exposed and straddles the interface of the scaffold and transport domains^18,19,22^, potentially precluding the need for hairpin-coordinated gating in DASS transporters. In this study, we have probed the substrate-dependent accessibility of several residues in the re-entrant hairpin loops of VcINDY; M157 and T154 in HP1, and S381, L384, and V388 in HP2 (Fig. 8A). Interestingly, we observe different effects of coupling ion and substrate binding within the same hairpin. As with other tested positions, T154 and S381 are mostly sensitive to the presence of Na^+^, whereas L384, V388, and in particular M157 exhibit considerable sensitivity to the presence of succinate (Fig. 5C). While further experimental work is required to provide a full picture of the dynamics of VcINDY’s re-entrant hairpins, these results suggest discrete effects of the coupling ion and substrate on the conformation of the hairpins.

**Figure 8.**
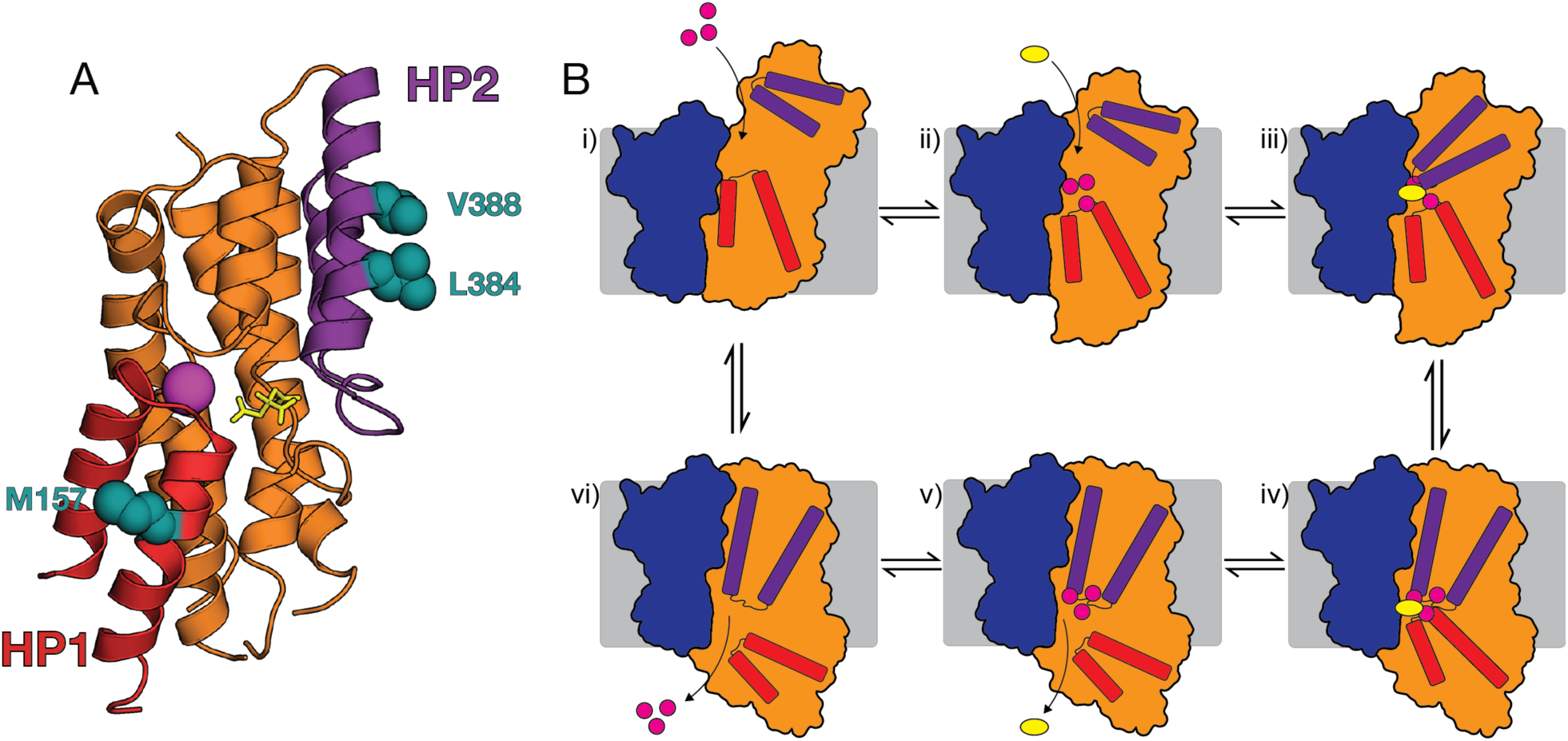
Position of succinate-sensitive hairpin residues and a hypothetical mechanism of VcINDY. **A)** Section of the X-ray structure of VcINDY’s transport domain highlighting HP1 (red), HP2 (purple) and the three positions particularly sensitive to the presence of succinate (teal). The substrate (citrate, yellow sticks) and Na^+^ ion (purple sphere) are shown. **B)** Hypothetical mechanism of VcINDY. (i) Outward-open VcINDY binds Na^+^ ions, which stabilises an open-intermediate state (ii); succinate binds, which stabilises the closed state of HP2 (iii), allowing the protein to transition to an inward-closed state (iv); HP1 opens (v); coupling ions and substrate are released into the cytoplasm (v & vi); empty transporter transitions back to the OFS to restart the cycle. Colour scheme is the same as A).

Analogous to the situation with Glt_Ph_ and its homologues, binding of Na^+^ is likely required for high affinity binding of succinate, and binding of the succinate molecule is required to stabilise the “closed” conformation of the re-entrant loop, which allows the elevator-like conformational change to take place^23,43-46^. The suggestion that the re-entrant hairpins undergo conformational changes in response to succinate binding is strengthened by the observations made during a series of solvent accessibility studies on the eukaryotic DASS transporter, NaDC1^47,48^. In concordance with our work presented here, the binding of Na^+^ and succinate had substantial and separable effects on the solvent accessibility of residues in the arm of re-entrant hairpin 2 in NaDC1 (in the same region and S381, L384 and V388 in VcINDY), suggestive of discrete conformational changes upon each Na^+^ and succinate binding event^47,48^. In contrast to our observations with VcINDY, the HP2 re-entrant loop residues become more accessible in the presence of Na^+^ alone, and then less accessible upon addition of succinate^47^. The differences between these two systems may be explained by the experimental set-up; NaDC1 was probed in the lipid bilayer, in the presence of gradients and a membrane potential, and in a background with 10 native cysteines present, whereas, in our study, each VcINDY variant was detergent-solubilised and contained only one cysteine residue. Cysteine accessibility studies on another bacterial DASS family member, SdcS from *Staphylococcus aureus*, revealed that D329 (F291 in VcINDY), which is predicted to be on the outward façade of the protein, is accessible in the presence of Na^+^, but not in the absence^49^. However, the equivalent position of VcINDY was not tested in this study. Curiously, the same SdcS study, N108 (N94 in VcINDY) was shown to be accessible from both the external and cytoplasmic sides of the membrane. However, in both the IFS crystal structure and the repeat-swapped model of VcINDY, there is dense protein blocking access to this site from the cytoplasmic side of the membrane^49^. This perhaps suggest that the mechanism of SdcS is considerably different to the current structural model based on VcINDY, or that the conformational changes during transport are *substantially* more extreme than currently thought. Nevertheless, these studies combined suggest that there are discrete conformational changes upon binding Na^+^ and succinate to DASS transporters. Indeed, solvent accessibility studies on NaDC1 prior to the elucidation of the structure of VcINDY revealed that the accessibility changes in HP2 were temperature insensitive, indicative that the modulation in accessibility was not due a *large* conformational change, but perhaps due to blockade of these possible binding site residues by succinate^47^. However, our current structural understanding of the DASS family reveals that these residues are in the arm of HP2 and do not form part of the binding site. Therefore, these observed accessibility changes of HP2 residues could be explained by subtle, local conformational changes of the hairpin in response to succinate binding. More structural information is required to fully realise the DASS transporter mechanism, in particular, a structure of the *apo* state of VcINDY will be especially illuminating.

### Potential implications to the mechanism of VcINDY

While further work is needed to fully realise the dynamics of VcINDY, our data are consistent with VcINDY having a complex mechanism that consists of multiple large-scale and local conformational changes as described by the following hypothetical mechanistic scheme (Fig. 8B). For simplicity, we will start this transport cycle with VcINDY in its open OFS. However, as VcINDY is a secondary-active transporter, it has the ability to work in the reverse direction depending on the direction and magnitude of the electrochemical gradients. In the open OFS conformation (Fig. 8B, state i), VcINDY is initially ligand free and the overall architecture resembles the structure described by the repeat swapped model^22^. One or more Na^+^ ions bind, inducing a shift in the transport domain conformation into an intermediate state in which the majority of sites tested in this study are obscured to some extent (Fig. 8B, state ii). However, HP2 remains “open” allowing succinate to still access the binding site. Binding of succinate stabilises the “closed” state of HP2 (Fig. 8B, state iii), allowing the transport domain to move into the closed IFS (Fig. 8B, state iv). HP1 opens (Fig. 8B, state v), releasing the substrates into the cytoplasm (Fig. 8B, state v and vi). HP1 closes and the ligand-free transport domain can translocate to form the OFS and restart the cycle.

### Substrate binding appears to be ordered and low affinity

In this study, we have identified a position in the arm of HP1, M157, whose accessibility is insensitive to the addition of Na^+^, but undergoes substantial accessibility changes in the presence of succinate (Fig. 5C and 6A). Probing this mutant in more detail, we discovered that the accessibility effects were only induced by the addition of compounds known to be transportable by VcINDY, for example, succinate, malate and fumarate, demonstrating specific binding was required (Fig. 7). In addition, we did not observe any changes in accessibility upon addition of succinate in the absence of Na^+^, demonstrating that Na^+^ must bind first before succinate can bind, as has been suggested in previous studies on DASS transporters^50^. This ordered binding is consistent with other Na^+^-driven elevator transporters. The archetypal elevator transporter, Glt_Ph_, transports aspartate coupled to the co-transport of 3 Na^+^ ions^51,52^, with 2 Na^+^ ions binding first to the *apo* transporter to ‘prime’ the binding site before the aspartate and final Na^+^ bind^46,53^, allowing the hairpin to close and the OFS-to-IFS transition to occur. Our data suggest a similar ordered binding mechanism for VcINDY. However, more detailed analysis of the coupling ion and substrate binding is required to illuminate this process further. When titrating succinate and monitoring the dose-responsive change in modification of VcINDYM157C^IFS^, we only observed substantial shifts in the modification efficiency once the succinate concentration reached ∼100 µM (Fig. 6B). This concentration is surprisingly high considering VcINDY’s *K*_m_ for succinate transport in proteoliposomes in the presence of a similar Na^+^ concentration is 1 µM^17^. To put this in the context of other elevator-like transporters; Glt_Ph_ also has *K*_m_ for transport of ∼1 µM^54,55^, but has a *K*_d_ of 100 nM^46^. Since VcINDY and Glt_Ph_ have a similar mechanism and identical *K*_m_s, it is not unreasonable to predict they would have similar *K*_d_s. However, if this were the case, we would expect VcINDY’s binding site to be saturated at 100 µM; the fact that the we continue to observe decreases in the modification rate with increasing succinate concentration demonstrates that it is not. These results indicate that in these experimental conditions VcINDY has a low affinity for substrate. Transporters with *K*_m_s lower than their *K*_d_ are rarely observed, one notable example being CaiT, which has a *K*_m_ of ∼80 µM and *K*_d_ of ∼3 mM^56^. As is the case with CaiT, VcINDY’s apparent low binding affinity may be due to the protein containing more than one functionally significant substrate binding site, as has been suggested for other DASS transporters^57^. Alternatively, this apparent low affinity may be due to VcINDY being in a detergent-solubilised state. Further functional analysis in the presence of a lipid bilayer are needed to resolve this issue.

We performed this study on detergent-solubilised protein in order to make the results directly comparable to the crystal structure of VcINDY, which was crystallised in its detergent solubilised form. In addition, we selected our panel of single cysteine mutants based on the crystal structure and repeat-swapped model of VcINDY in order to report on the these particular conformations.

Using lipid bilayer mimetics, such as detergents, may produce different results to what is seen in a membrane environment, where there is an absence of lateral pressure, other membrane proteins, specific protein:lipid interactions and electrochemical gradients. Indeed, single molecule FRET (smFRET) experiments have demonstrated that Glt_Ph_ has appreciably different dynamic behaviour in detergent micelles versus lipid bilayer^40,58^. However, site-directed alkylation studies on other transport proteins have directly shown that alkylation rates are similar in detergent compared to those measured in lipid bilayer^59^, giving us confidence that the qualitative differences we observe in the presence and absence of substrates are mechanistically relevant. The elevator-like mechanism requires large-scale conformational changes, which is almost certainly affected by the lipid composition (both lipid head group and hydrocarbon chain length and saturation). The dynamics and overall transport rate of the only other well characterised elevator-like transport, Glt_Ph_, is influenced by the presence and composition of the lipid environment and molecular dynamic simulations have recently shown large scale bilayer deformation can be induced by elevator-like transporter conformational changes^40,58,60-63^. So, aside from any specific protein:lipid interactions that may be influencing VcINDY’s function, the physical properties (e.g. flexibility) of the bilayer, which is dictated by the lipid composition, will clearly have a strong effect on its mechanism. How the lipid environment and electrochemical gradients across bilayers influences protein dynamics and substrate-dependent conformational changes of VcINDY is of great interest and is the subject of ongoing studies in our lab.

## MATERIALS AND METHODS

### Molecular biology and cysteine mutants generation

All single cysteine variants were generated in a previously characterised cysteine-free background in which all three native cysteines had been substituted for serine^17^. Substitutions were made with a Quikchange II site-directed mutagenesis kit (Agilent Technologies). Expression plasmids were fully sequenced to ensure that the desired codon substitution occurred and no unwanted secondary mutations were introduced.

### Expression of VcINDY

Over the course of this work, we modified the expression protocol of VcINDY three times in an effort to maximise its yield, which led to an increase in yield from 0.2 mg/L culture to >5 mg/L culture for wildtype VcINDY. Changes in expression levels of VcINDY did not affect the quality of the protein produced, the PEGylation efficiency for each variant, nor the transport activity of each variant (data not shown). In all three protocols, VcINDY and its variants were expressed in-frame with an N-terminal decahistidine tag form a modified pET vector^64^. VcINDY was initially expressed as described previously^17,18^. Briefly, BL21-AI (Invitrogen) cells harbouring the expression vector were grown in LB supplemented with 50 µg/ml kanamycin and incubated at 37°C until it reached an A_600_ of 0.8. Cells were rapidly cooled in an ice bath for 20 min, at which point expression was induced by addition of 10 µM IPTG and 6.6 mM L-arabinose. The cells were incubated at 19°C and grown for approximately 16 hours before being harvested and resuspended in Lysis Buffer (50 mM Tris, pH 7.5, 200 mM NaCl, and 10% (v/v) glycerol). Following disappointing yields via the above method, we adopted the MemStar method described by Drew and co-workers^65^. Briefly, in this protocol, Lemo21 (DE3) (New England Biolabs) cells harbouring the expression plasmid were grown in PASM-5052 media^66^, which was supplemented with 50 µg/ml kanamycin and 25 µg/ml chloramphenicol and incubated at 37°C until it reached an A_600_ of 0.5. At this point, expression was further induced by the addition of 0.4 mM IPTG and the cells were incubated for approximately 16 hours at 25°C before being harvested and resuspended in Lysis Buffer.

The following small modifications to this protocol increased the yield of VcINDY approximately 10-fold; the PASM-5052 media was exchanged for the MDA-5052 and the kanamycin concentration was increased to 100 µg/ml. These changes likely led to an increase in VcINDY yield because of better maintenance of the kanamycin resistant expression plasmid in the Lemo21 (DE3) cells. Lemo21 (DE3) (and other BL21 derivatives) grow robustly in high levels of kanamycin in high phosphate media, such as PASM-5052^66^, whereas, in MDA-5052 media, which contains half the phosphate concentration, Lemo21 are again sensitive, making the antibiotic selection of expression plasmid-containing cells more effective.

### Purification of VcINDY

VcINDY was purified as detailed previously^17^. Briefly, resuspended cells were lysed by sonication, the lysate was clarified by centrifugation at 20000 xg for 20 min, and the membrane fraction was isolated by ultracentrifugation at 200000 xg for 2 hours. Membrane vesicles were resuspended in Purification Buffer (50 mM Tris, pH 8, 100 mM NaCl, 5% (v/v) glycerol). For the purification of cysteine containing variants of VcINDY, the Purification Buffer was supplemented with 0.5 mM tris-(2-carboxyethyl)phosphine (TCEP) to keep the cysteines in a reduced state. 0.5 mM TCEP was added to all purification buffers. VcINDY was solubilised by incubating the vesicles with 19.6 mM n-dodecyl-β-D-maltopyranoside (DDM, Glycon) for 1 hour at 4°C. Non-solubilised material was removed by ultracentrifugation and the soluble fraction was incubated with Talon metal affinity resin (Takara Bio) for 16 hours at 4°C. Loosely bound contaminants were eluted from the resin and the detergent exchanged by 2 rounds of washing using Purification Buffer supplemented with 1.96 mM DDM and 10 mM imidazole, followed by Purification Buffer containing 5.4 mM n-decyl-β-D-maltopyranoside (DM, Glycon) and 20 mM imidazole. Protein was eluted by incubating the resin with Purification Buffer supplemented with 5.4 mM DM and 10 µg/ml trypsin for 1 hour at 4°C. The purified protein was concentrated and stored at −80°C. All cysteine-containing protein were stored in the presence of TCEP to keep the cysteines reduced.

### Protein reconstitution

Protein was functionally reconstituted as detailed previously^22^. 25-100 µg of DM-solubilised and purified protein was diluted to 2 ml in Reconstitution Buffer (25 mM Tris, pH 8, 100 mM NaCl, 5% glycerol and 3% DM) and mixed with 400 µl 20 mg/ml *E. coli* polar lipids (Avanti Polar Lipids). This mixture of protein/lipid was incubated on ice for 10 min followed by rapid dilution into 65 ml of Inside Buffer (20 mM Tris, pH 7.5, 1 mM NaCl, 199 mM KCl, and 1 mM DTT). The resultant proteoliposomes were collected by ultracentrifugation, resuspended in Inside Buffer to a concentration of 8 mg/ml lipid, freeze thawed thrice, and stored at −80°C.

### *In vitro* transport assays

For transport assays, the required amount of proteoliposomes were thawed, extruded 11 times through a 400 nm filter, collected by ultracentrifugation, and resuspended to a final concentration of 80 mg/ml lipid. Transport assays were performed by rapidly mixing the prepared proteoliposomes with Reaction Buffer (20 mM Tris, pH 7.5, 100 mM NaCl, 100 mM KCl, 1 µM valinomycin, and 1 µM [^3^H]-succinate (American Radiolabelled Chemicals). At frequent timepoints, samples were collected from the transport reaction and quenched by addition of ice cold Quench Buffer (20 mM Tris, pH 7.5, 200 mM ChCl). Proteoliposomes and the accumulated [^3^H]-succinate were collected by rapid filtration through 200 nm nitrocellulose filters (Millipore). The filters were washed with 3 ml of Quench Buffer, dissolved in FilterCount liquid scintillation cocktail (PerkinElmer) and the accumulated [^3^H]-succinate was counted using a Hidex 300SL Liquid Scintillation Counter.

### PEGylation timecourse

To perform the PEGylation timecourse, detergent solubilised protein was thawed and buffer exchanged using Zeba Spin Desalting Columns (Thermo Fisher Scientific) to remove the 0.5 mM TCEP and exchange the protein into PEGylation Buffer (50 mM Tris, pH 7, 5.4 mM DM, 5% (v/v) glycerol). The substrate-free apo sample was formulated by mixing protein solution with 1 M KCl and PEGylation Buffer to generate a final protein concentration of 10 µM and a final KCl concentration of 150 mM. The “Na^+^ alone” sample was formulated in the same way except KCl was substituted for NaCl. The same approach was used for the “Na^+^ + succinate” samples except that succinate was added to a final concentration of 1 mM. For the “succinate alone” sample, protein was mixed with 150 mM KCl and 1 mM succinate. The protein/substrate mixtures were incubated for at 10 minutes at room temperature, at which point the PEGylation reaction was started by addition of methoxypolyethylene glycol maleimide 5K (mPEG5K) (Sigma Aldrich). Samples were collected at various timepoints and the reaction was terminated by addition of SDS-PAGE sample buffer containing 100 mM methyl methanethiosulfonate (MMTS) (Sigma Aldrich). The PEGylation reaction samples were analysed using non-reducing polyacrylamide gels, which were stained with Coomassie Brilliant Blue dye to visualise the protein.

### Densitometric analysis

The intensities of the bands corresponding to the unmodified and PEGylated VcINDY protein bands were quantified using ImageJ software^67,68^. For each timepoint, the modification efficiency was calculated using the following equation;

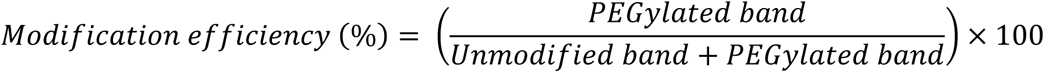

For the final data sets, replicates (n≥3) of the modification efficiency for each timepoints were averaged and the standard error of the mean (SEM) was calculated. Where indicated, statistical significance was examined using unpaired t-tests.

## Acknowledgements

We thank L. Forrest, V. Leone, and G. Fitzgerald for helpful discussions. We thank A. Quigley for discussions and suggestions regarding maximising the yield of VcINDY. This work was supported by the University of Kent Graduate Teaching Assistantship (GTA) programme, a Wellcome Trust Seed Award in Science (210121/Z/18/Z) awarded to CM, and the Division of Intramural Research of the US National Institutes of Health, National Institute of Neurological Disorders and Stroke. We would like to dedicate this work to the memory of H. Ronald Kaback.

## Author contributions

C.M. and J.A.M. conceived project; C.M., C.D.D.S. and M.J.S. performed research and analysed data; C.M. wrote the manuscript; all authors reviewed the manuscript.

## Additional information

The authors declare no competing interests.

## Supplementary Figures

**Supplementary Figure 1.**
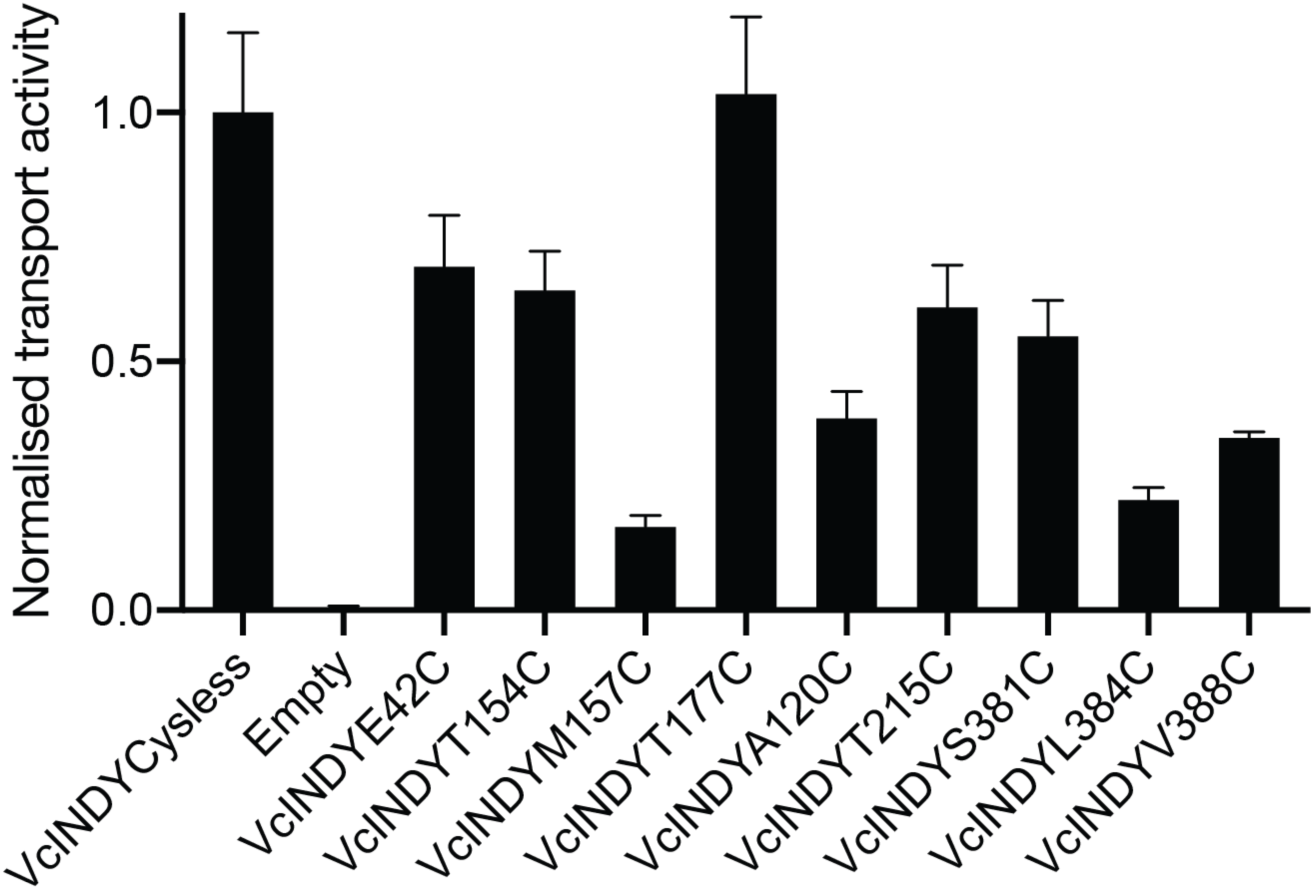
Transport activity of all 9 single cysteine variants. Initial rates of [^3^H]-succinate transport into proteoliposomes containing cysteine-free VcINDY (VcINDYcysless) or one of the 9 single cysteine variants, normalised to VcINDYcysless activity. Background levels of [^3^H]-succinate accumulation were determined using a protein-free liposome control (“Empty”). Data points are the average of triplicate datasets and the error bars represent SEM. This experiment was performed at least twice for each mutant with the same result.

**Supplementary Figure 2.**
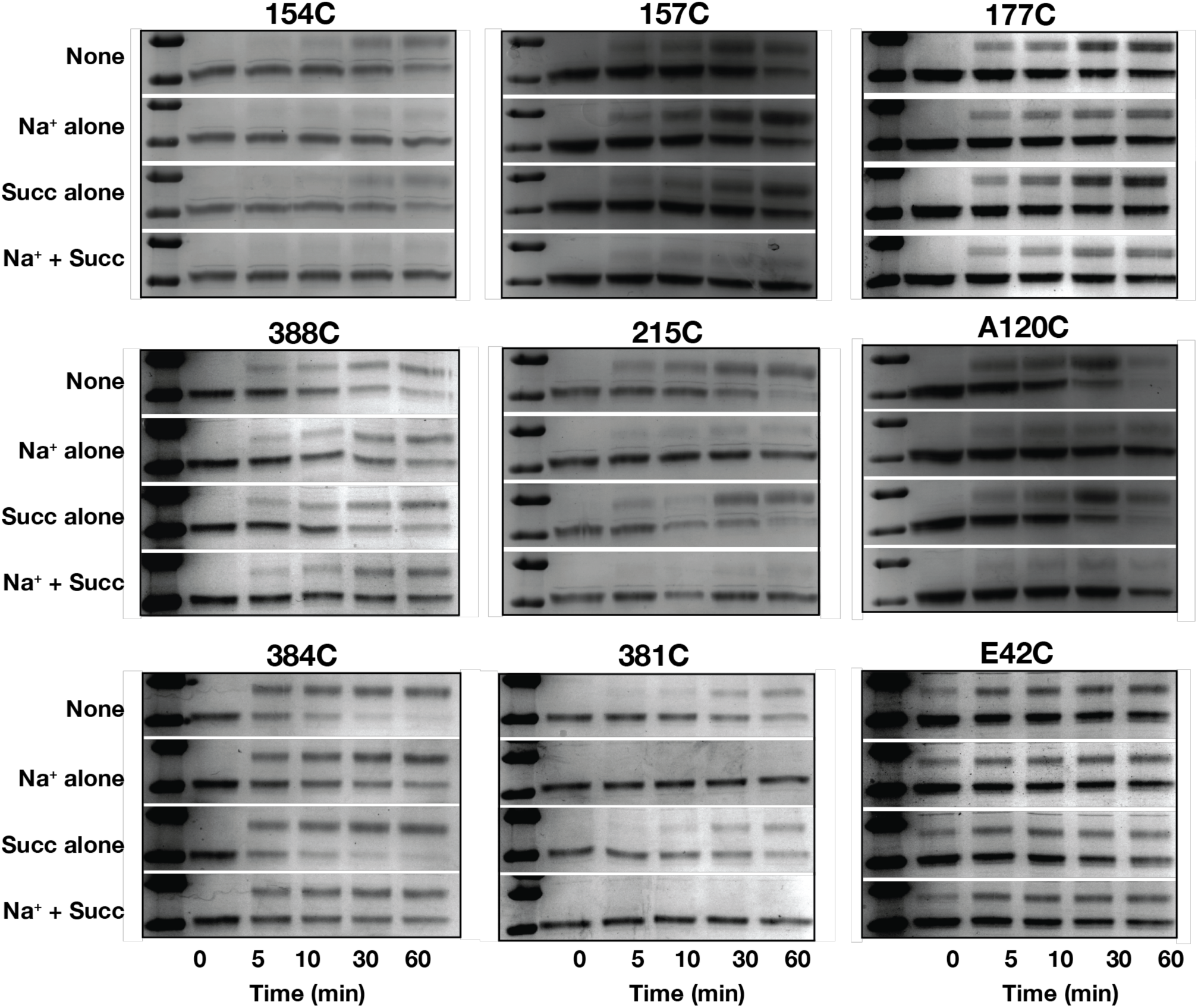
SDS-PAGE analysis of PEGylation rate of all 9 single cysteine variants. Representative gels of the modification timecourse of each single cysteine variant tested under each substrate condition; no substrates (150 mM K^+^), Na^+^ alone, succinate alone (in 150 mM K^+^), and Na^+^ and succinate.

**Supplementary Figure 3.**
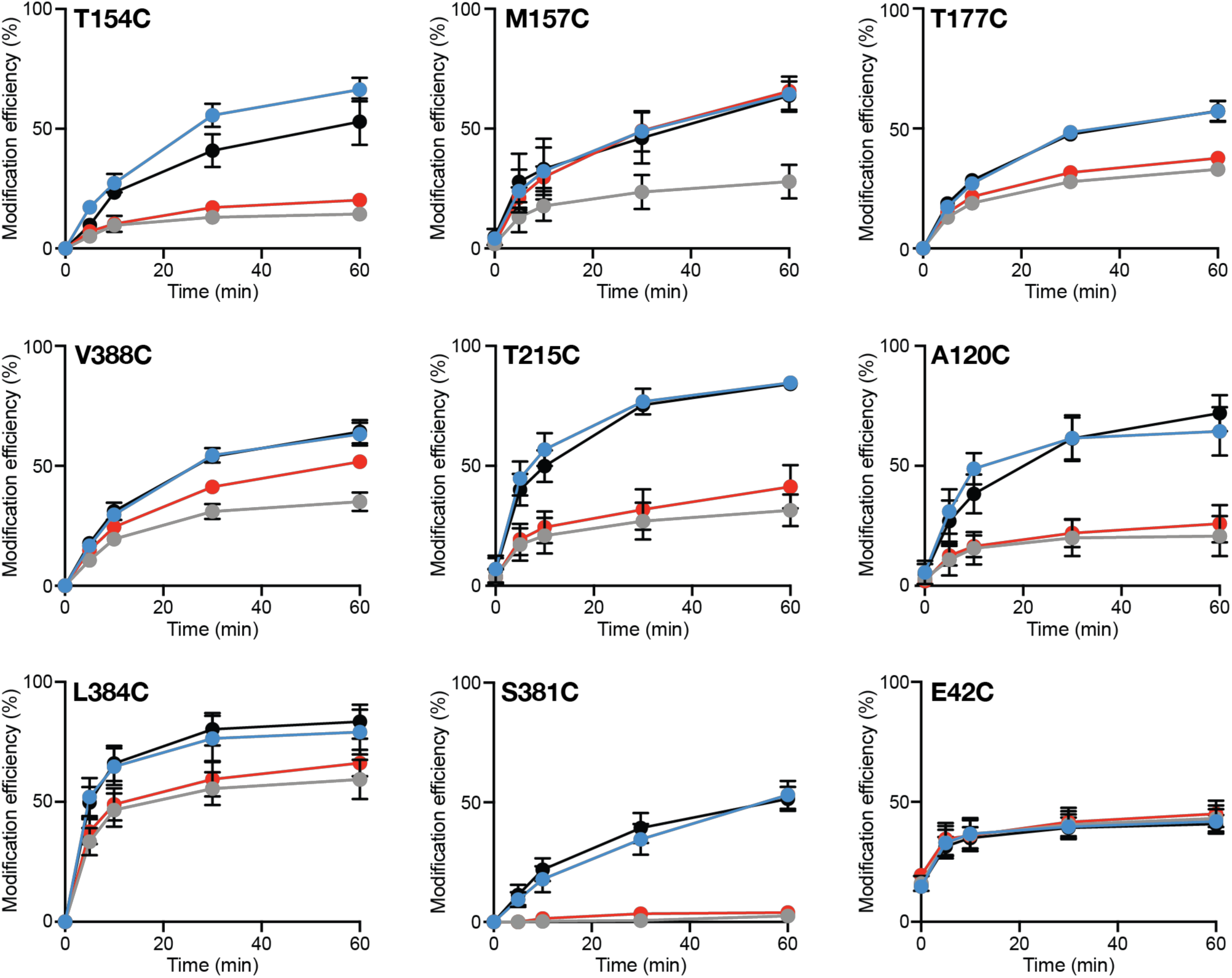
Densitometric analysis of PEGylation rate of all 9 single cysteine variants. Proportion of each single cysteine mutant modified at each timepoint under different substrate conditions; no substrate (blue data), succinate alone (black data), Na^+^ alone (red data), and Na^+^ and succinate (grey data). Data points are the average of triplicate datasets and the error bars represent SEM. This experiment was performed on at least 3 separate occasions for each mutant with the same result.

**Supplementary Figure 4.**
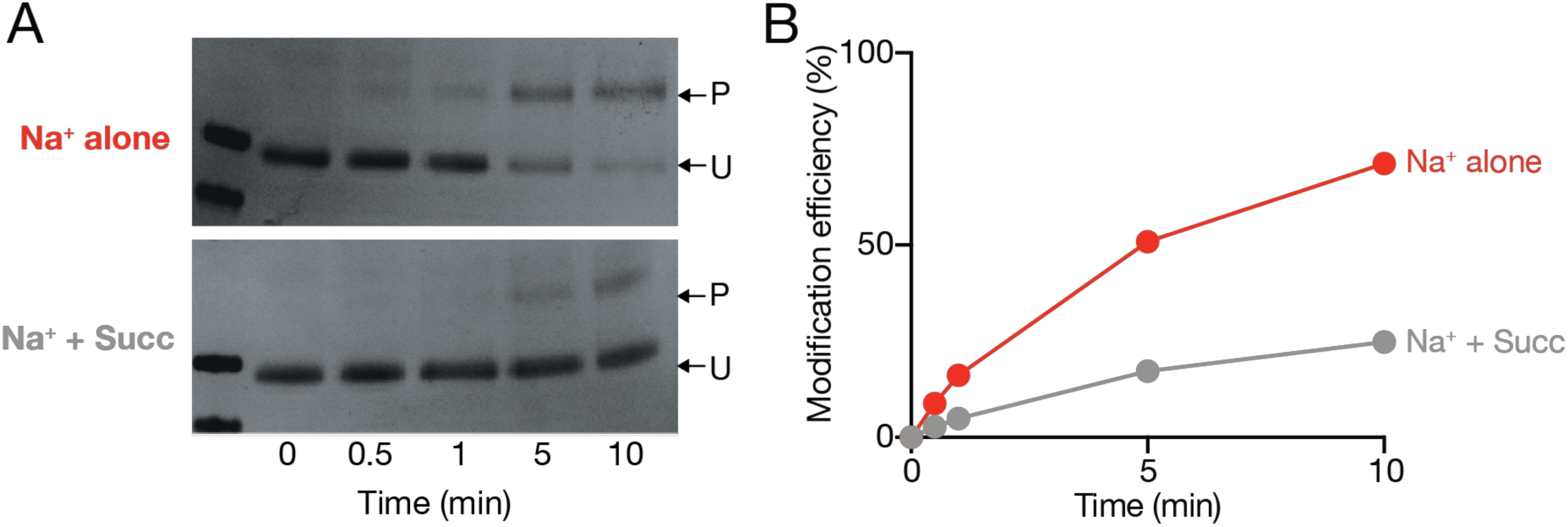
Substrate induced accessibility changes of VcINDYM157C^IFS^ measured by modification by MTS-PEG5K. **(A)** SDS-PAGE gel of the MTS-PEG5K modification timecourse of VcINDYM157C^IFS^. The PEGylated protein bands (P) and unmodified protein bands (U) are indicated by arrows. **(B)** Densitometric analysis of (A). Proportion (%) of VcINDYM157C^IFS^ modified at each timepoint in the presence of 150 mM Na^+^ alone (red data) or in the presence of 150 mM Na^+^ and 1 mM succinate (grey data).

